# Microstructural–functional coupling as a multimodal biomarker of emotional state transitions in bipolar disorder

**DOI:** 10.1101/2025.11.14.688566

**Authors:** Lei Wei, Dongsheng Wang, Shaoyang Fang, He Wang

**Author notes:** Lei Wei and Dongsheng Wang contributed equally to this work as the co-first authors. He Wang, Lei Wei, contributed equally to this work as co-corresponding authors.

## Abstract

Bipolar disorder (BD) is characterized by dynamic transitions between depressive and manic states, yet the neural mechanisms underlying these state shifts remain unclear. Here, we developed an integrative framework to quantify the microstructural–functional coupling (MFC) of the brain, which captures the distributional similarity between voxel-wise diffusion and functional features within each cortical region. Using multimodal MRI data from 72 BD patients and 65 matched healthy controls, we observed a global increase of MFC of BD, which indicates a de-similarity of brain microstructure and function. Partial least squares correlation (PLSC) analysis revealed that the principal latent component linking MFC to behavioral scores explained 67.2% of the variance, with the strongest contributions from the control and somatosensory networks. Mediation analysis demonstrated that regional MFC influenced anxiety indirectly through depressive symptoms, supporting a sequential affective transition from depression to anxiety. Moreover, macroscale gradient analysis showed that the principal MFC gradient was significantly correlated with neurotransmitter receptor distributions, including α4β2, 5-HT1B, and H3. Together, these findings highlight MFC as a multimodal biomarker sensitive to emotional state transitions in BD and suggest its potential as a bridge between microstructural alterations, functional dynamics, and neurotransmitter systems.

## Introduction

Bipolar disorder (BD) is a prevalent neuropsychiatric illness that often emerges during young adulthood. Patients with BD experience recurrent episodes of mania and depression, reflecting dynamic transitions between distinct brain states [1]. In most cases, patients initially present with depressive episodes and later develop manic symptoms [2]. This dynamic transition not only increases the difficulty of early diagnosis but also leaves the underlying mechanisms of such state shifts poorly understood [3].

Cortical microstructure has been shown to be sensitive to various mental disorders. Using free-water imaging, patients with BD have exhibited significantly increased free-water content across widespread gray matter regions, suggesting potential neuroinflammatory processes and metabolic abnormalities [4]. In addition, cortical mean diffusivity (MD) has been suggested as an important biomarker towards Alzheimer’s disease (AD) [5, 6]. Cortical fractional anisotropy (FA) has also been suggested to serve as a potential biomarker for detecting or characterizing neurological disorders [7]. Generally, cortical microstructure has shown the ability to distinguish patients from healthy controls (HC), and also provided biological explain to the cortical morphological or functional alterations of mental disorder patients [8]. Although cortical microstructural measures derived from dMRI can capture subtle alterations in local cytoarchitecture, they may not directly reflect changes in behavior or cognition. Functional MRI (fMRI) is commonly regarded as an effective approach to link brain activity with altered behavioral performance. The power spectrum–based amplitude of low-frequency fluctuations (ALFF) and the regional homogeneity (ReHo) of BOLD time series have been widely used in neuropsychiatric research [9-12]. More specifically, reduced functional connectivity within the default mode network (DMN) has been associated with impaired emotional and cognitive regulation in patients with BD [13, 14], whereas aberrant coupling within the limbic system has been linked to affective instability [15]. In major depressive disorder (MDD), previous studies have reported an abnormal pattern of network expansion, suggesting altered large-scale functional organization in depression [16].

The activity of cortical fMRI signals primarily depends on the underlying regional cytoarchitecture [17]. This structure–function dependence can also be represented at the network level. For example, the DMN shows a distinctive microstructural organization, and its heterogeneous cytoarchitecture may account for its unique functional properties [18]. Recent network-based analyses have further highlighted the role of microstructural features in shaping functional connectivity, revealing a macroscale correspondence between microstructure and function [19]. Collectively, these findings suggest that local coupling between microstructural and functional properties provides a mechanistic basis for large-scale brain connectivity. Structural–functional coupling has thus been increasingly investigated through network analyses from various perspectives. Since dMRI and fMRI capture different aspects of neuropsychiatric processes, their combination may provide deeper insights into the mechanisms underlying mental disorders. Recent studies have proposed constructing voxel- or vertex-wise morphological feature matrices across brain regions and computing interregional similarities at the matrix level [20]. This approach disregards the spatial arrangement of voxels or vertices within individual regions and instead focuses on the overall distributional similarity between regional pairs, providing a distinct nonlinear perspective on regional similarity.

Using this framework, our study aims to integrate microstructural and functional metrics to gain a more comprehensive understanding of how multimodal regional phenotypes contribute to neuropsychiatric disorders. Microstructural features provide a basis for assessing neuroinflammation and cytoarchitectural alterations, whereas functional metrics offer direct insights into externally observable behavioral manifestations. On the one hand, microstructural measures capture the initial biological alterations, while functional metrics provide a deeper understanding of dynamic functional organization. Together, these two aspects may form the mechanistic link underlying state transitions in neuropsychiatric disorders.

## Methods

### Data acquisition

A total 72 BP patients and 65 matched healthy controls were enrolled from the Psychiatric Department of the First Affiliated Hospital, School of Medicine, Zhejiang University.With informed consent from each subject, MR scans were performed on a 3T MR imaging system(The Signa HDx 3.0T of General Electric Healthcare)with an 8-channel phased array head coil. Subjects were placed in a supine position, and the coil was filled with a sponge pad to keep the head stationary during MRI scanning. In order to improve subjects’ comfort, earplugs were used to lighten the noise during the examination. During the resting-state functional MRI scanning, all subjects were asked to close their eyes, rest quietly, stay awake, and avoid systematic thinking.

Whole-brain high structural MRI data were collected using a three-dimensional magnetization-prepared rapid gradient echo sequence with the following imaging parameters: repetition time (TR)□=□7084 ms; echo time (TE)□=□2.8 ms; inversion time□=□600 ms; matrix size□=□256□×□256; 156 slices; slice thickness□=□1 mm; voxel size□=□1.0□×□1.0□×□1.0 mm^3^; number of averages =1; field of view□=□240 mm; and flip angle□=□8°. Whole-brain resting-state fMRI data were collected using a T2*-weighted gradient-echo-planar imaging sequence with these parameters: TR□=□1800 ms; TE□=□30 ms; slice thickness□=□4 mm; spacing between slices = 4.8; matrix size□=□64□×□64; voxel size□=□3.75 ×□3.75□×□4 mm^3^; number of averages =1; 180 time points; field of view□=□240 mm; and flip angle□=□90°. Spin-echo(SE) echo planar imaging (EPI) sequence was used to acquire diffusion MRI (dMRI) with following parameters: TR□=□11000 ms; TE□=□89.4 ms; slices = 44; slice thickness = 3 mm; matrix size = 128 × 128; field of view□=□240 mm; number of averages =2; flip angle□=□90°;b-values=1000s/mm2; number of diffusion directions=25.

### Neuropsychiatric testing

AssessmentDepression severity was evaluated using both clinician-rated and self-reported instruments. The clinician-administered 17-item Hamilton Depression Rating Scale (HAMD) and the Montgomery–Åsberg Depression Rating Scale (MADRS) were employed. The HAMD, scored from 0 to 52, was interpreted using established cut-offs for severity. The MADRS, scored from 0 to 60, was included for its sensitivity to changes. Self-reported depressive symptoms were measured using the Patient Health Questionnaire-9 (PHQ-9), where a score of ≥10 indicates a likely depressive episode.Manic symptoms were assessed using the clinician-administered Young Mania Rating Scale (YMRS), with scores ranging from 0 to 60; a score of ≥12 was used to indicate clinically significant mania. All clinician-administered scales (HAMD, MADRS, YMRS) were conducted by trained personnel who established inter-rater reliability.

### Data preprocessing

All T1-weighted images were processed using FreeSurfer v7.4 [21] to generate cortical and subcortical segmentations. Structural images were further registered to the MNI152 template space using the ANTs SyN algorithm (https://github.com/ANTsX/ANTs).

Functional MRI data underwent standard preprocessing, including slice-timing correction and realignment within the individual space. The preprocessed data were then normalized to the MNI152 space with a 3 mm isotropic voxel size. To minimize partial-volume effects from different tissue types, spatial smoothing with a 4 mm full width at half maximum (FWHM) Gaussian kernel was applied only within the gray matter mask. Subsequently, white matter and cerebrospinal fluid (CSF) signals, and 24 head motion parameters (24P) were regressed out. The residual time series were temporally band-pass filtered within 0.01–0.1 Hz. ALFF and ReHo maps were then computed using a standard fMRI analysis pipeline [22]. Dynamic ALFF maps were estimated using a sliding-window approach with a window length of 30 time points and a one-timepoint step. Finally, voxel-wise temporal signal-to-noise ratio (tSNR) maps were generated to assess the quality of the fMRI signal.

Diffusion MRI preprocessing was performed using MRtrix3 [23]. The raw diffusion data were first denoised and corrected for Gibbs ringing artifacts. After EDDY current and motion correction, the dMRI images were normalized to the MNI152 space with a 2 mm isotropic voxel size. To minimize CSF contamination, a single shell freewater elimination model was applied [24, 25]. Freewater–corrected FA, MD, axial diffusivity (AD), and radial diffusivity (RD) maps were subsequently computed for each subject. All resulting diffusion maps were then resampled to match the spatial resolution and template space of the fMRI images.

### Within-regional microstructure and functional activity coupling

For each cortical and subcortical region, fMRI-based and dMRI-based features were extracted separately. To quantify within-regional functional connectivity, voxel-wise correlation matrices were computed using all voxels within each brain region. The mean connectivity strength of each voxel was derived as a voxel-level feature, resulting in an M × N feature matrix for each region, where M denotes the number of voxels within the region and N denotes the number of fMRI-derived features. In our current study, we used the Schaefer 100 cortical atlas to parcellate the brain

For diffusion MRI, a similar procedure was applied. Tensor-derived metrics including FA, MD, AD, and RD were combined to form voxel-wise diffusion feature matrices. Voxels containing more than 80% free water were excluded to avoid CSF contamination, yielding an M × P feature matrix for each region, where M is the number of valid voxels and P is the number of diffusion-derived features.

The resulting fMRI- and dMRI-based feature matrices represented the distributions of functional activity and microstructural properties within each region. To quantify the cross-modal similarity between these two representations, we developed a voxel-level alignment and comparison approach. Each feature matrix was first standardized across voxels using z-score normalization and reduced to a common dimensionality using principal component analysis (PCA). In the current study, the first three principal components were retained for both functional and microstructural feature matrices.

Because PCA yields components with arbitrary orientations between datasets, we applied an iterative closest point (ICP) alignment to register the two feature distributions into a common coordinate space. At each iteration, the algorithm identified, for each sample in the functional feature matrix, its nearest neighbor in the microstructural matrix and then estimated the optimal rigid orthogonal transformation that minimized the distance between the two feature sets. This process was repeated several times to reduce potential bias introduced by PCA decomposition and ensure a more robust cross-modal alignment. In our current study, we repeated 5 times for ICP alignment.

After alignment, the similarity between the two feature distributions was quantified by computing the symmetrized k-nearest-neighbor Kullback–Leibler (KNN-KL) divergence between the functional and microstructural matrices [20]. A smaller divergence indicates a higher degree of similarity between functional and microstructural feature representations within that region.

### Network based functional-microstructural coupling

For each group, we constructed a group-wise affinity matrix by computing pairwise MFC between brain regions. Age, sex and TIV were regressed out before the correlation matrix computation. The resulting group-level affinity matrices were used to extract macroscale gradient by using BrainSpace toolbox [26]. Specifically, we applied the diffusion map embedding approach to derive gradient components that capture the principal axes of functional connectivity variation across the cortex. Using the neuromaps library [27], we further assessed the spatial similarity between the network gradient maps and 19 neurotransmitters to compute the correlation with MFC principle gradient [28].Spin permutation was used to determine the significance of each correlation. The resulting p-value was further corrected by FDR correction.

### Partial least regression analysis

In the current analysis, 6 behavioral measurements were included: the Patient Health Questionnaire-9 (PHQ-9), Hamilton Depression Rating Scale (HAMD), Hamilton Anxiety Rating Scale (HAMA), Young Mania Rating Scale (YMRS), Generalized Anxiety Disorder-7 (GAD-7), Montgomery–Åsberg Depression Rating Scale (MADRS). Partial least squares correlation (PLSC) was used to examine regional associations between microstructural–functional coupling and neuropsychiatric status. Age, sex, and total intracranial volume (TIV) were regressed out before PLSC analysis. The statistical significance of each latent component was assessed using 5,000 permutation tests, and the reliability of brain–behavior loadings was further estimated via bootstrap resampling. This approach enabled the evaluation of the robustness and stability of the decomposed brain–behavior relationships. The whole PLSC analysis was implemented by using myPLS toolbox [29].

### Statistical analysis

A linear regression model was used to compare the differences between BD patients and HC. Age, sex and TIV were considered as covariates for the comparison. For each brain region, we compute the group comparison significance and correct the p-value by using FDR correction. Mediation analysis was conducted using the Process5.0 R scripts. Age, sex, and TIV were included as covariates for mediation analysis.

## Results

### Group differences in microstructural–functional coupling between BD and HC

We first applied the microstructural–functional coupling (MFC) metric to compare patients with bipolar disorder (BD) and healthy controls (HC). The results revealed that group differences were primarily localized within the default mode network (DMN). Specifically, coupling values in the BD group were moderately decreased in the right posterior cingulate cortex (PCC) but increased in the left prefrontal cortex (PFC) and right orbitofrontal cortex (OFC). After FDR correction, the coupling value in the left PFC remained significantly higher in BD compared with HC (t = 3.67, FDRp = 0.03; Figure 2).

**Figure 1.**
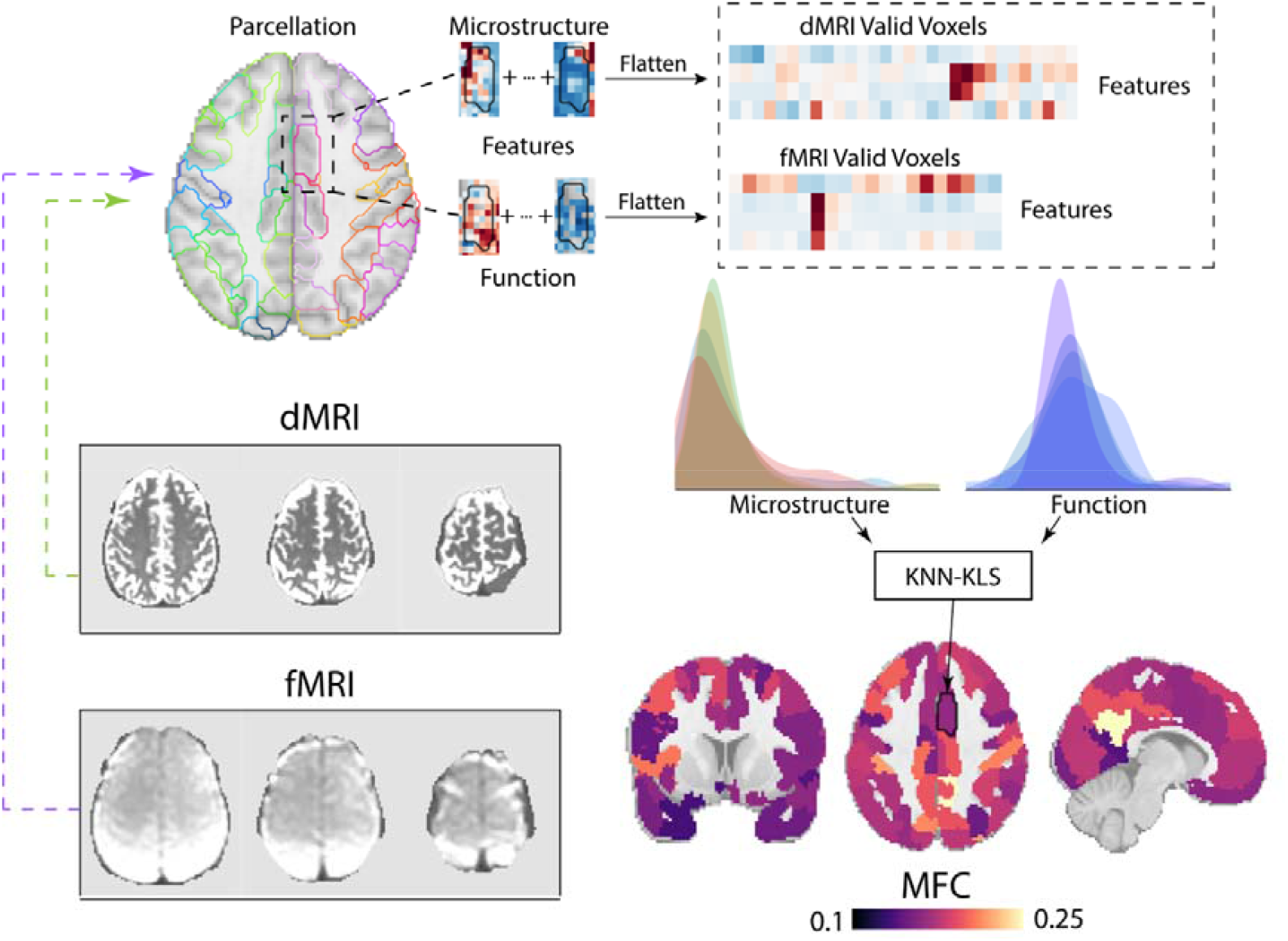
Estimation of MFC.

**Figure 2.**
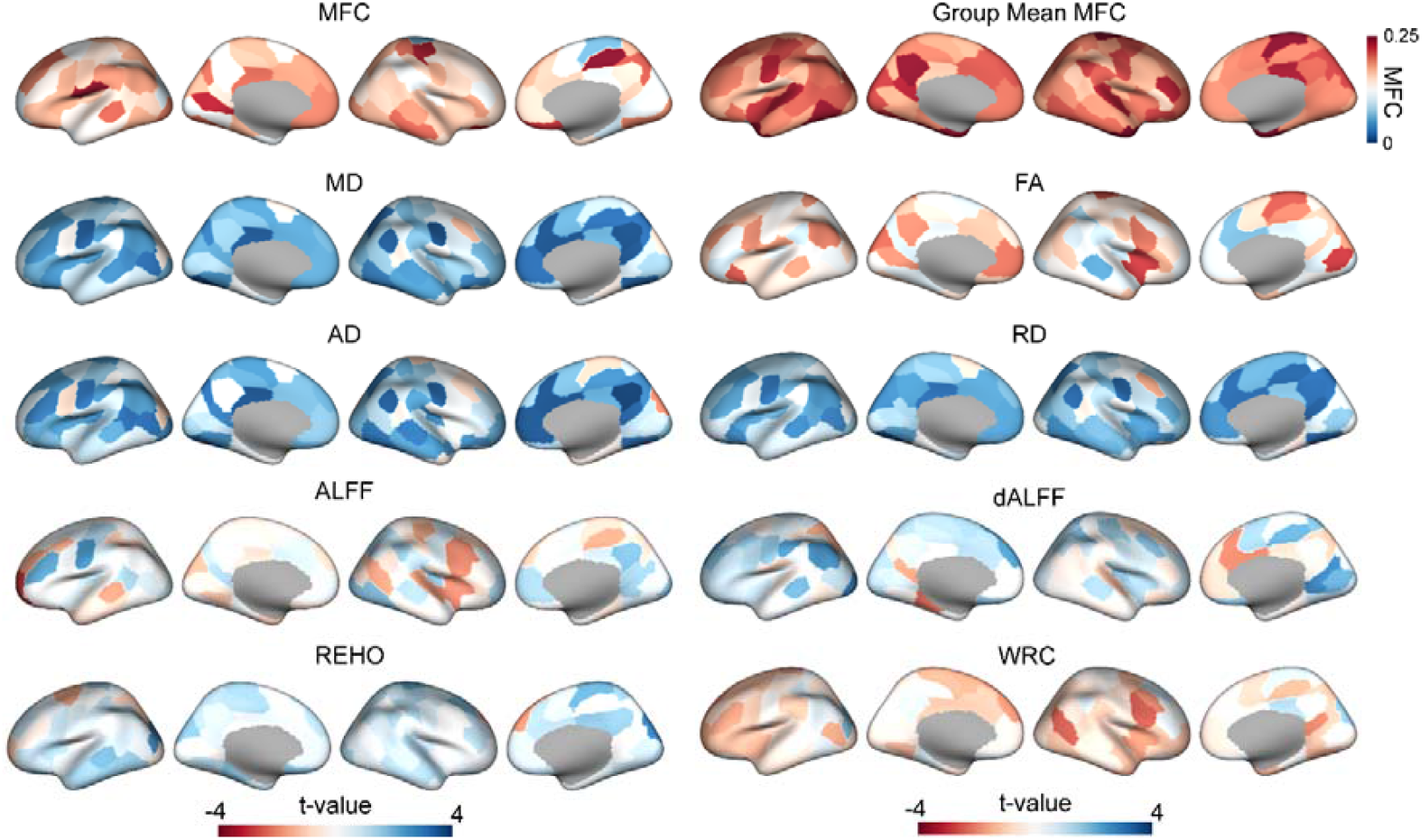
Group comparison of the functional and microstructural map between BD and HC.

In addition, we examined group differences in regional functional and microstructural measures separately between BD and HC (Figure 2). In the current dataset, parcellation-based comparisons of functional measures alone showed limited sensitivity to BD-related abnormalities, whereas microstructural indices exhibited more spatially specific alterations. Notably, axial diffusivity (AD), mean diffusivity (MD), and radial diffusivity (RD) demonstrated significant reductions in diffusivity around the parietal lobe, particularly within regions belonging to the DMN, dorsal attention network (DAN), and control network. However, distinct spatial patterns were observed among these diffusivity metrics. An apparent increase in fractional anisotropy (FA) was found in the PCC of BD patients, although this difference did not survive FDR correction. Moreover, the PCC also exhibited subtle but noticeable group differences in AD, MD, and RD values.

Group-level patterns of microstructural–functional coupling (MFC) and diffusion metrics between patients with bipolar disorder (BD) and healthy controls (HC). The averaged MFC map across all participants exhibited a medial–lateral gradient across the whole brain. The BD group showed a significantly increased MFC value in the left prefrontal cortex (t = 3.67, p = 0.0003). No significant group differences were observed for other functional metrics. Group comparisons of diffusion measures (AD, MD, and RD) revealed similar spatial patterns, with distinct alterations around the posterior cingulate cortex (PCC).

Regions within the bilateral somatosensory, dorsal attention, and control networks showed significantly decreased diffusivity in BD (Table S1). No significant group differences were found in FA.

MFC, microstructural–functional coupling; WRS, average nodal strength of within-regional voxel connectome; ALFF, amplitude of low-frequency fluctuations; dALFF, dynamic ALFF; ReHo, regional homogeneity; FA, fractional anisotropy; AD, axial diffusivity; RD, radial diffusivity; MD, mean diffusivity; BD, bipolar disorder; HC, healthy controls.

### Latent decomposition of the brain–behavior covariance

The PLSC analysis was adopted to further decompose the relationship between MFC and bipolar behavior. Mania and depression behavior scores estimated by PHQ9, HAMD, GAD-7, HAMA, MADRS, and YAMARS were used for this analysis. The first latent variable (LV1) explained 67.2% variance of the behavior. The results were significant after 5,000 permutations (p_perm = 0.012, Figure 3A). The spatial pattern of LV1 exhibited a clear lateral–medial organization across the whole brain. The BSR map shows that the regions within control and somatosensory network contribute mostly to the brain-behavior covariance.

**Figure 3.**
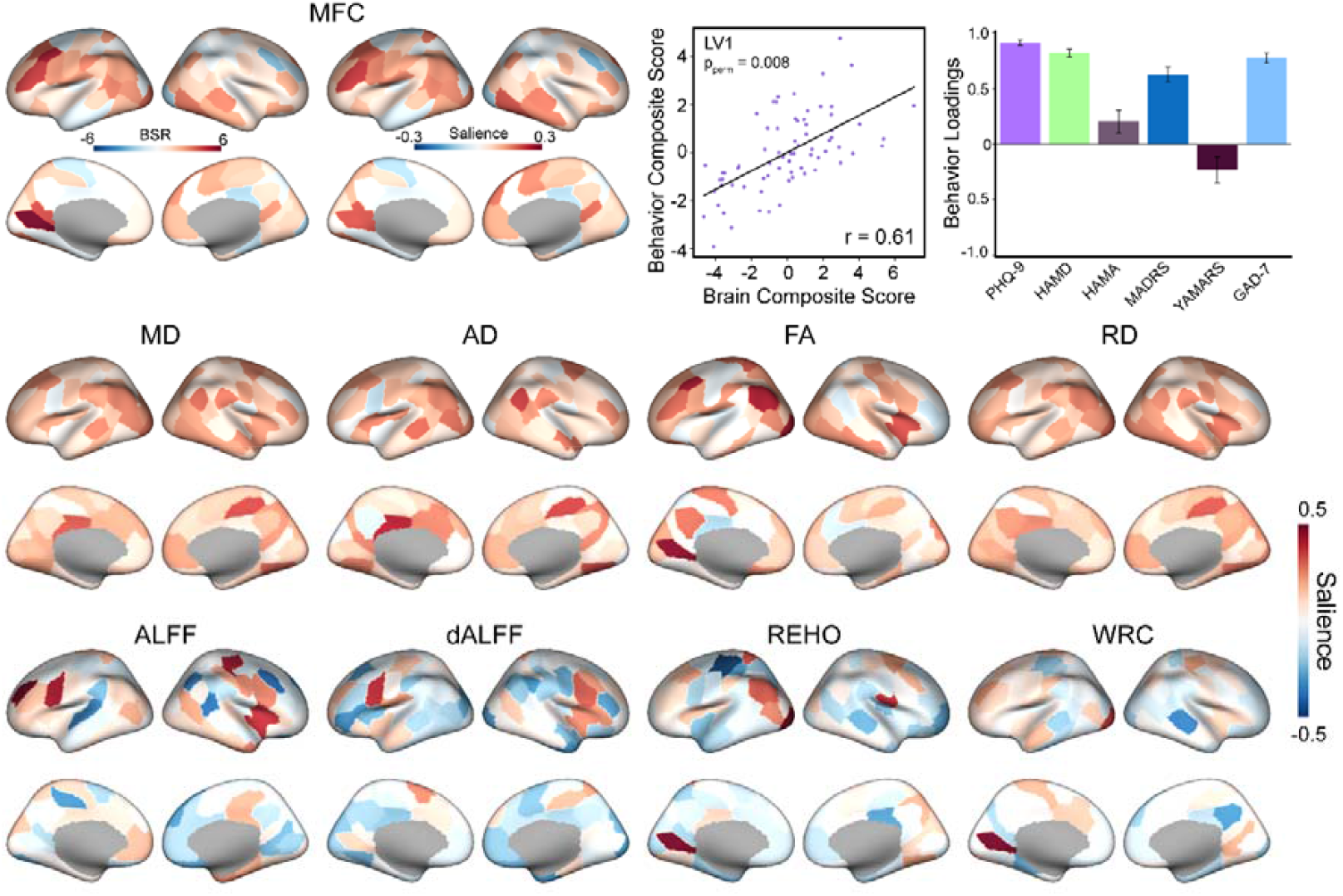
Partial least square correlation results of MFC and BD behavior. A Whole brain bootstrap ratio and brain salience of the MFC. The LV1 explained 67.2% of the brain–behavior covariance and reached statistical significance after 5000 permutations (p = 0.012). B,C depicts the modality-specific bootstrap ratio maps obtained from the PLSC solution for each of the eight MRI modalities (functional and microstructural).

We further applied PLSC analyses to all eight functional and microstructural MRI features. No significant LV was found from all these 8 features. The bootstrap ratio (BSR) maps corresponding to the latent variables that explained the largest proportion of variance are shown in Figure 3B–C. Overall, the microstructural maps exhibited weaker sensitivity to bipolar behavioral measures, whereas the functional maps showed stronger associations with the behavioral latent variable (Figure 4).

**Figure 4.**
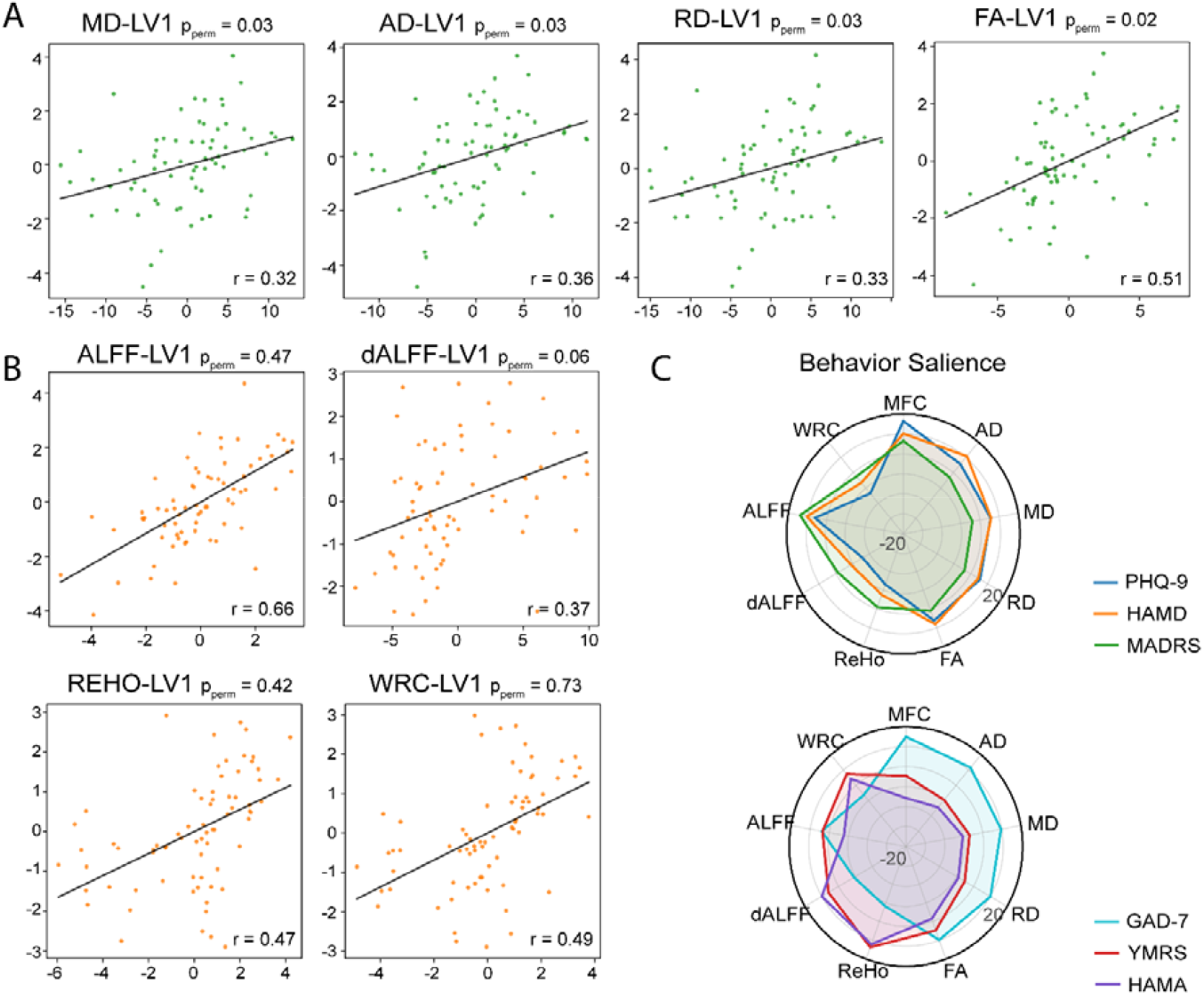
A. Correlations between the LV1 from PLSC analyses across the four microstructural features. B. Correlations between the first LV1 from PLSC analyses across the four functional features. C. Radar plot showing the behavioral salience of LV1 for each of the eight features.

### MFC Influences Anxiety Indirectly Through Depressive Symptoms

We first extracted the first principal components (PC1) of depression and anxiety scales separately. A mediation analysis was then conducted with depression PC1 as the predictor, anxiety PC1 as the mediator, and the YMRS-based mania score as the outcome. The results revealed a significant indirect effect of anxiety on the relationship between depression and mania (ab = 2.92, 95% CI [1.63, 5.02]). Depression exhibited a significant negative direct effect on mania (c′ = −5.06, 95% CI [−6.85, −3.27]). Because the direct and indirect effects were in opposite directions, this pattern reflects a suppression mediation, indicating that higher depression scores reduce mania directly but also increase mania indirectly through elevated anxiety levels. To further validate these findings, we repeated the mediation analysis using the PHQ-9 and GAD-7 scores separately, as the PLSC results indicated that MFC was most strongly correlated with these two scales (Figure S1). This analysis yielded a similar suppression mediation pattern (ab = 0.78, 95% CI: [0.33, 1.50]).

We then incorporated the first principal component of whole□brain MFC (MFC PC1) into a serial mediation model (Figure 5). The analysis revealed a significant serial indirect effect through depression and anxiety linking MFC to mania (a×b×c = 0.25, 95% CI [0.04, 0.60]). This suggests that reduced MFC integrity may indirectly influence manic symptoms via increased depressive and anxious symptoms. Furthermore, we tested the brain regions with bootstrap□ratio (BSR) values greater than 4 identified in the PLSC analysis. Four regions included two subregions in the left DMN, one in the left somatomotor network, and one in the left DAN. All four regions exhibited significant serial indirect effects through regional MFC, depression, anxiety, mania, consistent with the whole□brain MFC result. We presented one region in DMN (a×b×c = 0.25, 95% CI [0.04, 0.60]) and one in DAN in Figure 5, the other two regions were presented in Figure S2.

**Figure 5.**
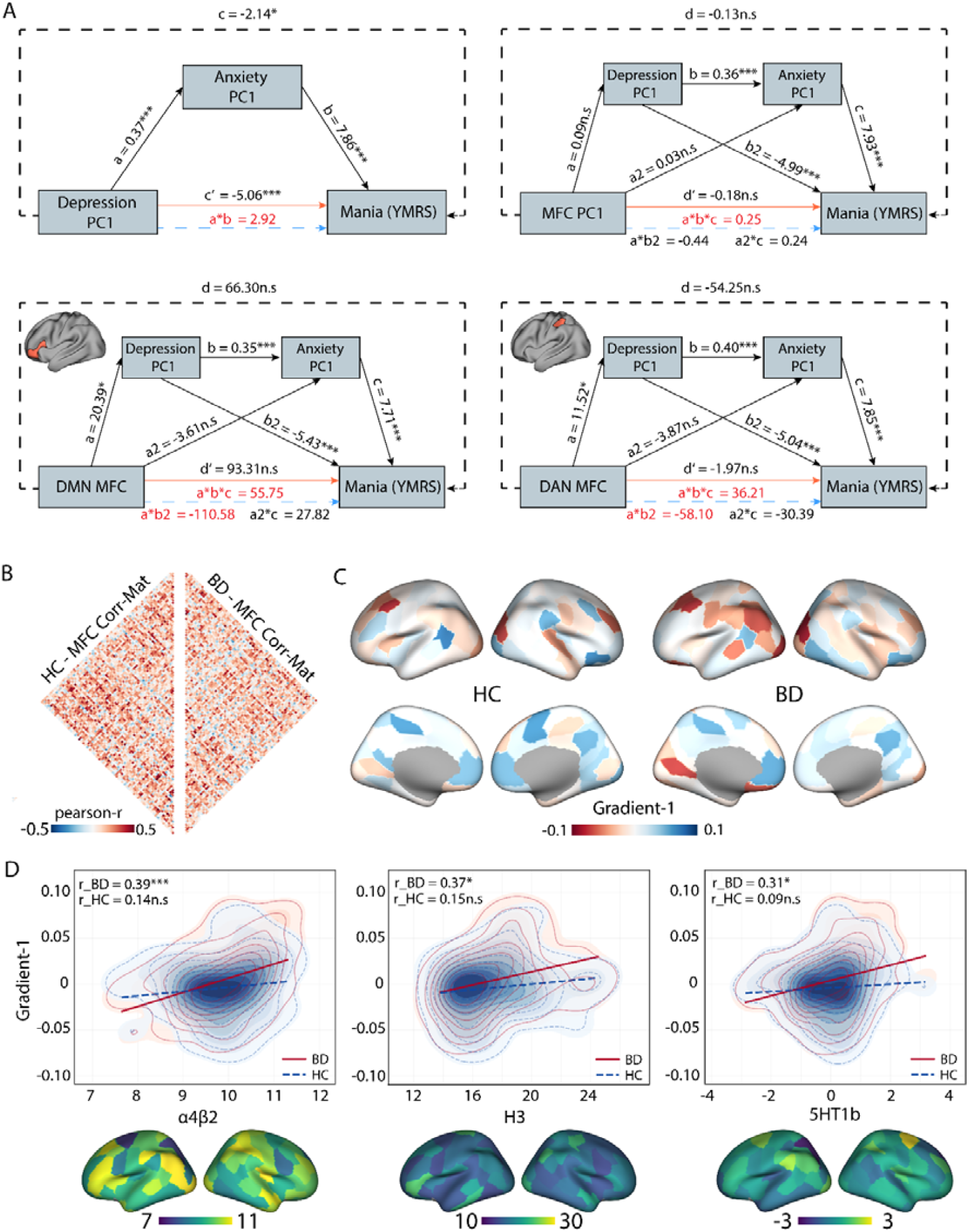
A. Results of the serial mediation analysis. Indirect effects highlighted in red indicate 95% confidence intervals that do not include zero. The blue dash line represents the indirect effect; the orange dash line represents the direct effect. B. Group-wise MFC correlation matrix. C. The first gradient decomposed by Brainspace of HC and BD respectively. D. Correlation between the first gradients and neurotransmitter maps. The first gradient of BD group showed significant correlated with the α4β2 (r = 0.39, p_spin < 0.001), H3 (r = 0.37, p_spin = 0.04), and 5HT1b (r = 0.31, p_spin = 0.01) after FDR correction. ***: p<0.001; *: p<0.05.

### Macroscale gradient of MFC across the whole brain associated with neurotransmitter distribution

Group-level whole-brain MFC similarity matrices were computed separately for the HC and BD groups (Figure 5.B). Network gradients were then generated from the combined matrix by using joint embedding approach. The proportion of variance explained by each gradient component was calculated (Figure S1). The first gradient, which accounted for the largest proportion of variance (14.8%), was used for subsequent analyses. For all 19 neurotransmitter systems, we found that the principal gradient in the BD group showed significant positive correlations with the acetylcholine receptor α4β2 (r = 0.39, p_spin < 0.001), the serotonin receptor 5-HT1B (r = 0.31, p_spin = 0.01), and the histamine receptor H3 (r = 0.37, p_spin = 0.04). No significant correlations were observed between the MFC principal gradient and any neurotransmitter receptor map (Figure 5C, D).

## Discussion

In this study, we integrated multiple microstructural and functional features within each brain region using a distributional similarity framework. Our results suggest that the combined feature, termed MFC, provides a more comprehensive perspective than analyzing individual microstructural or functional metrics alone. Given the distinct biological roles of brain microstructure and function, we further propose that MFC is sensitive to state transitions in BD patients, particularly from depressive to manic phases.

Group-level analyses revealed that microstructural features exhibited greater sensitivity to differences between patients and HCs, while the behavior PLSC analysis showed that the functional features were more sensitive to the abnormal behavior changes. Recent research indicates that the cytoarchitectural organization of regions such as the default mode network (DMN) underlies cross-regional and hierarchical signal integration within the brain [18]. Evidence from perfusion and calibrated fMRI studies suggests that changes in oxygen metabolism within the DMN modulate global patterns of functional connectivity [30]. More specifically, the heterogeneity of cortical cytoarchitecture and its influence on brain function profoundly affect energy transitions within the brain at the macroscale. This relationship further highlights the critical role of microstructure–function coupling in facilitating state transitions of the brain [31]. Since BD usually experiences frequent brain state transitions, we observed a general increase in MFC across the whole brain, indicating a decoupling between microstructural and functional organization. Furthermore, we suggested that the global increase of MFC might mainly contribute by the similar pattern of cortical diffusivity. Unlike AD, decreased diffusivity might cause by more complicate reasons. Previous studies indicate that the increase of diffusivity was found during the brain development stage, relating to the myelin process and neuronal dendrite increase, which limited the interstitial water diffusion [32]. In addition, increased of cortical diffusivity has also been found within early stage AD patients, suggesting the early stage inflammation response and cortical morphometry changes [33]. In our current results, we suggest that the observed decrease in diffusivity should be interpreted in conjunction with increased fundamental functional activity. The elevated ALFF and WRC values indicate enhanced within-region communication, which may reflect a state of regional hyperactivation rather than simple functional suppression. Overall, we propose that reduced cortical diffusivity provides the microstructural basis for cortical hyperactivity. This phenomenon may reflect increased dendritic density or an early compensatory response within the cortex.

According to our PLSC analysis, functional features presented a higher sensitivity to the external behavior scores. It should be noted that in our current study, we attempted to categorize the behavioral features of BD into two main dimensions: activation and inhibition. Anxiety often co-occurs with depression, and both are generally considered negative affective traits [34]. Previous study also suggest that anxiety frequently emerges during the course of depression and may be associated with manic traits or mixed affective states [35]. Within the framework of emotional continuity [36, 37], the progression from negative–low arousal to negative–high arousal and finally to positive/irritable–high arousal states could be interpreted through alterations in MFC. According to the results of the mediation analysis, a general triangular relationship was observed among depression, anxiety, and mania. Moreover, we found that regions within the dorsal attention network (DAN) and the DMN exhibited a significant mediation pathway through regional MFC, linking depression, anxiety, and mania. Notably, these regions also showed higher BSR values in the PLSC analysis. Increased MFC within the DAN and DMN was consistent with the reported alterations in large-scale intrinsic networks in BD [38]. Moreover, the strong correlation between the group-level MFC principal gradient and 5-HT1b receptor density suggests that MFC may serve as a potential intermediary link between the serotonergic system and the abnormal integration of the sensorimotor and higher order nonmotor networks, such as the DMN [38, 39]. The other two receptor that related to MFC principle gradients, including H3 and α4β2 also associated with the mood regulation [40, 41]. Given these properties, we propose that the MFC is highly sensitive to emotional arousal. It may act as an initiating node that responds to and drives the sequential transformation of emotions along a specific trajectory. This interpretation is consistent with our mediation results showing a chained mediating effect from depression to anxiety to mania through regional MFC activity. However, the MFC itself did not directly mediate the effect of anxiety on mania, suggesting that its influence may primarily lie in initiating, rather than maintaining, this emotional transition.

Nevertheless, several limitations should be acknowledged. First, we did not incorporate morphological characteristics. Although morphology was not the primary focus of the present study, it could provide complementary information to microstructural features and allow for a more comprehensive interpretation of MFC alterations. Second, our findings were limited to bipolar disorder; future studies including other psychiatric populations will be essential to determine the generalizability and specificity of MFC as a cross-diagnostic biomarker. In this study, we incorporated within-regional connectivity (WRC) as part of the functional features. WRC was used to characterize voxel-level connections within each region, thereby capturing the local coherence of neural signals. However, it should be noted that WRC is not a fully robust metric, as it may be affected by local signal autocorrelation. The inclusion of WRC was therefore intended primarily to provide an estimate of the local integration strength of regional signals, which could partially correspond to microstructural alterations. Indeed, we found that WRC exhibited meaningful correlations with behavioral features, further enhancing the behavioral interpretability of the MFC framework.

In summary, by integrating microstructural and functional features, we proposed a novel psychopathological biomarker — the microstructural–functional coupling (MFC). MFC combines the advantages of both modalities, providing not only a robust ability to differentiate patients from healthy controls but also a finer-grained interpretation of individual behavioral characteristics. In patients with bipolar disorder, MFC successfully identified specific brain regions that represent the directional transitions of emotional states.

